# Nonlinear distributed sensing of light patterns leads to perceptual distortions in plants

**DOI:** 10.64898/2026.03.31.715481

**Authors:** Ahron Kempinski, Amir Porat, Mathieu Rivière, Yasmine Meroz

**Author notes:** equal contribution (Ahron Kempinski, Amir Porat).

## Abstract

Many organisms lack centralized sensory-processing systems, navigating complex environments through local integration of spatially distributed stimuli (e.g., chemotaxis of cells, bacteria, or growing neurons). Here we propose the first general physical and geometric framework to describe how such distributed sensing translates into integrated directional responses. We study plants, multicellular decentralized systems that grow towards light (phototropism), which can come from multiple directions and at different intensities. We develop a model in which light is sensed locally on the shoot circumference, transduced nonlinearly, and integrated vectorially; the model is informed and validated by unilateral and bilateral lighting, and out-of-plane illumination experiments on sunflower seedlings. We show that seedlings respond to the vectorial sum of transduced signals rather than to the physical sum of incident light, which can create systematic deviations between maximal physical illumination and growth direction, akin to optical illusions. This framework further predicts that symmetrical, opposing cues cancel each other out, which we validate experimentally using a weaker symmetry-breaking light source.

## Introduction

To navigate complex environments, organisms must extract directional information from spatially structured stimuli. In many animals, sensory inputs are collected by spatially distributed receptor arrays, such as the lateral line system in fish or mechanoreceptors in the skin (1, 2), and subsequently integrated by centralized nervous systems. In other organisms, however, both sensing and integration are distributed across extended surfaces, so that coherent responses emerge directly from the combination of local signals. This distributed strategy is observed across biological scales and sensory modalities: unicellular bacteria and amoebae integrate chemical gradients through receptors distributed over their membrane (3, 4); motile immune cells (5) and growing axons (6, 7) combine chemical and mechanical cues across their surface; certain cyanobacteria perform phototaxis using spatially distributed photoreceptors shaped by the geometry of the cell body (8); and large slime molds and fungal networks likewise coordinate spatially resolved signals without centralized control (9–11).

Researchers have long sought to understand how organisms that rely on distributed sensory integration combine multiple local directional cues into a single behavioral output. Numerous mathematical models have been developed for this purpose (3, 12–16). with most focusing on intracellular signaling dynamics and adaptation. Yet these models’ grounding in specific molecular implementations hinders them from providing a general physical and geometric understanding of distributed sensory integration, i.e., a framework linking nonlinear distributed sensing over extended surfaces to the integrated directional response. Such a framework is crucial for interpreting experimental observations in which organisms exposed to competing or complex gradients show responses that are nonlinear, hierarchical, or biased rather than simple reflections of the maximal physical stimulus (7, 16–18). Plants, multicellular organisms that lack a centralized processing system, provide a striking and tractable model system for exploring questions of distributed sensory integration. Plants survive and prosper by identifying favorable growth directions on the basis of stimuli such as light and gravity and redirecting growth accordingly; these processes are called tropisms (19–22). Macroscopic tropic responses are successfully described by theoretical models representing stimuli such as light and gravity as vectors (23–28). The molecular underpinnings of tropisms have also been extensively investigated. For example, in phototropism, growth in response to light, light is sensed primarily by phototropin receptors in the epidermis, leading to asymmetric growth across the organ cross-section (29, 30). Nevertheless, current understanding of tropisms remains insufficient for closing the gaps in our understanding of distributed sensory integration. Specifically, studies of the molecular pathways and macroscopic dynamics of phototropism have focused on responses to single light sources. Yet, natural environments expose plants to heterogeneous illumination from multiple directions (31). In particular, although phototropic responses do not always reflect simple maximization of light availability (32–35), it remains unclear how plants encode and integrate spatially distributed light cues, and how this internal representation is shaped by the geometry and nonlinear transduction of the sensory system.

Here we provide first insights into these questions by conducting controlled unilateral and bilateral lighting, and out-of-plane illumination experiments on sunflower seedlings (*Helianthus annuus*) and combining them with a three-dimensional theoretical framework for phototropism. We develop an experimentally informed model in which light is sensed locally on the shoot circumference (partially transmitted to the other side), transduced nonlinearly, and subsequently summed. We find that seedlings respond to the vectorial sum of transduced signals rather than to the physical sum of light stimuli. This mechanism produces systematic discrepancies between actual maximal illumination and perceived illumination (as reflected in growth direction), effectively generating *optical illusions* in plant phototropism. By making explicit the geometric origins of these effects, our results place plant tropic responses within a broader class of distributed sensory systems and provide a quantitative framework for directional integration in extended biological sensors.

## Results

### Physical properties of plant tissue inform theoretical model of spatial perception rules

Focusing on plant phototropism, we develop a conceptual framework describing how plants perceive and integrate multiple light signals coming from different directions in space. To do so, we build on a previously developed vectorial model describing the phototropic response of plant organs to a single light source (25, 37, 38). In broad strokes, the model approximates the shoot as a twistless and shearless slender rod, defining a variable *s* along the centerline, with *s* = 0 at the base and *s* = *L*(*t*) at the tip, equal to the length of the organ at time *t* (Fig. 1A). At every point *s* we define a local frame of reference using the Frenet-Serret framework (26), where 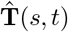 and 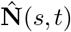 are the tangent and normal to the centerline, respectively, and the local curvature, which describes how much the organ is bent, follows 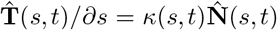, with the curvature vector defined as 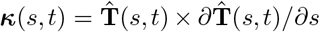 (inset of Fig. 1A). For the sake of clarity, from here on we drop the explicit dependence on *s* and *t*. Next, we describe external directional stimuli, such as light and gravity. Both are vector fields, and due to the minimal centerline description of the plant organ, they can be approximated by a single effective vector: a light source is described by its intensity *I* and direction **Î**, i.e. **I** = *I***Î**, and gravity is defined by the magnitude of acceleration *g* and its direction 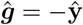, aligned with the lab frame of reference: 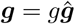. It is well established that only the components perpendicular to the centerline, marked **I**^⊥^ and ***g***^⊥^ (Fig. 1A), lead to changes in organ shape; that is, light or gravity parallel to the shoot do not cause tropic bending (24). The dynamics of tropic responses to these stimuli are described by:

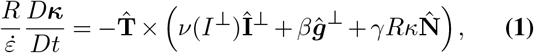

where the left-hand side describes the tropic response, manifested by the change of the organ’s curvature *κ*, or shape, over time. *R* is the radius, 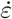 is the average growth rate, and we use a material derivative 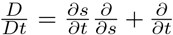, where 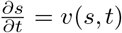 is the local velocity due to growth. The change in shape is driven by a differential growth rate across the plant cross section, which we define as a vector **Δ** across the organ circumference, depicted graphically in the inset of Fig. 1A (details in SM D). The growth gradient is in turn dictated by the sum of transduced stimuli; The prefactor in the phototropic term **Δ**_ph_ = *ν*(*I*^⊥^)**Î**^⊥^ is the transduced or *perceived* magnitude of the light intensity 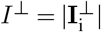 (24). The next term 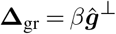 represents gravitropism (shoots grow upwards in response to gravity), with *β* the gravitropic sensitivity (23, 39) (independent of acceleration intensity (39)), and the last term represents proprioception or autotropism, with *γ* the proprioceptive sensitivity, and can be thought of as a restorative force, describing the tendency of plants to grow straight with no external stimuli.

**Fig. 1.**
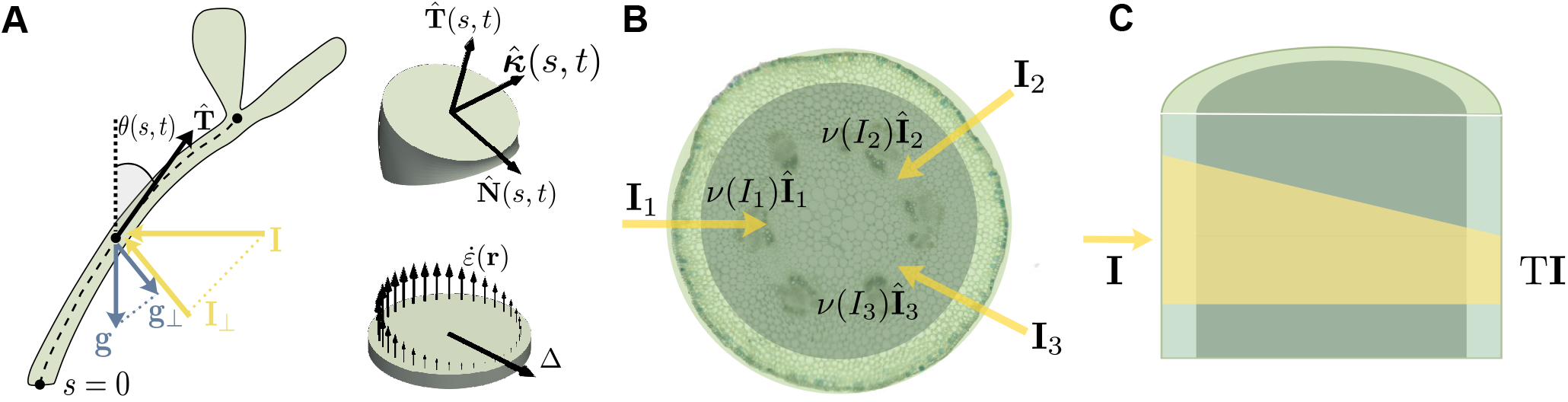
Geometric and physical basis of distributed light sensing and perception. **(A) Centerline representation of tropic dynamics.** The shoot is modeled as a slender growing rod parameterized by arc length *s*. External stimuli, including light **I** and gravity **g**, are vectors projected onto the local cross-sectional plane (**I**_⊥_, **g**_⊥_), driving curvature changes through differential growth across the organ. Tropic dynamics are restricted to the plane defined by the plant and the light source, and the shape of the plant is defined by the local angle *θ*(*s, t*) between the tangent 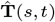 and the vertical. (top inset) At every point *s* we define a local frame of reference using the Frenet-Serret framework (26), where 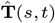 and 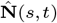 are the tangent and normal to the centerline, respectively, and 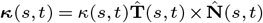 is the local curvature. (bottom inset) Differential growth, underpinning changes in curvature, is defined as the gradient of the growth rate vector field in the cross-section 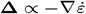 (details in SM D). **(B) Photoreceptors distributed around the shoot circumference**. Graphic representation of a typical cross-section of a plant shoot. Phototropin receptors reside primarily in the epidermis (highlighted), implying that incident light is sensed and transduced locally around the shoot circumference, before integration. **C. Light transmission through plant tissue**. A fraction T of incident light is transmitted to the opposite side of the shoot, producing a weaker counteracting signal. This transmission modifies the effective phototropic stimulus (36).

Building on this description, we now ask how plants represent and process spatial information about multiple light sources from different directions, **I**_i_, shaping their response. From a physics perspective, the total illumination is given by the vector sum ∑_i_ **I**_i_. The extensive experimental evidence corroborating Eq. 1 (23, 24, 40, 41), limited to a single light source, supports the fundamental idea that, in the context of tropisms, plants are able to encode directional stimuli as vectorial representations, and respond to a vectorial sum of one light source and gravity. Put together, these concepts suggest that plants respond to some form of vectorial sum of light stimuli.

To determine the vectorial summation law, we examine the geometry and optics of the light sensory system of a typical plant shoot. Photosensing occurs through photoreceptors called phototropins, mainly located in the outer layer of cells called the epidermis (represented graphically in Fig. 1B). When a shoot is exposed to multiple light sources from different directions, each source illuminates a different part of the shoot, and is therefore sensed and transduced locally on the circumference, following *ν*(*I*_i_)Î_i_ (Eq. 1), assuming uniform sensing restricted to the circumference. This suggests that plants may respond to the vectorial sum of the transduced signals, not the sum of the light signals as may be naively expected from an optics perspective.

Thus, for N light sources the phototropic signal in Eq. 1 is replaced with the expression:

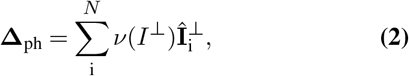

assuming different light directions 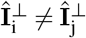.

Furthermore, light is partially transmitted through plant shoots (31, 36). Thus, a fraction of the light intensity, denoted by the transmission factor T, reaches the photoreceptors on the other side of the tissue, providing a weaker signal in the opposite direction of the original light stimulus (Fig. 1C). The phototropic term in Eq. 1 then takes the form

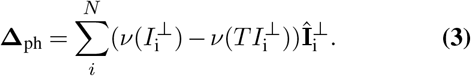

Here, we assume that the transmitted light remains confined to the cross-sectional plane, effectively modeling the shoot as a strongly disordered scattering medium, and neglect refractive effects (42). If two light sources are in the same or opposite directions, 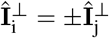, light intensities need to be summed accordingly before transduction, as detailed later on. In order to corroborate this model, we first estimate the phototropic transduction function *ν*(*I*), as detailed in the next section.

### Estimation of phototropic transduction function *ν*(*I*)

Sensory systems across diverse species generally transduce physical signals non-linearly (43, 44), typically following either Weber-Fechner’s logarithmic law (45), or Stevens’ power law (46). Changes in perception are then proportional to relative changes in stimulus intensity, rather than absolute changes, enabling organisms to efficiently process a wide range of stimulus intensities, from faint to overwhelming. An intuitive example is the non-linear human perception of loudness, reflected by the well-known logarithmic decibel (dB) scale used to measure sound intensity, where a 10dB increase is perceived as roughly double in loudness (47).

In plant phototropism, the perception of light is affected by factors such as fluence rate, internal structure, and previous exposure to light (etiolation) (36, 48–50). Building on previous observations (24) we assume the phototropic transduction function follows a power-law:

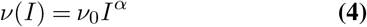

where *ν*_0_ is a prefactor, and *α* the power.

To quantify this function, we analyze the dependence of the magnitude of the phototropic response on light intensity.

We exposed sunflower seedlings, *Helianthus annuus*, to blue light from a single LED panel, for a range of light intensities spanning *I* ∈ [0.7, 7] W*/*m^2^ (Fig. S1). Tropic dynamics are restricted to a single plane defined by the plant and light source. We measured the tropic response of seedlings exposed to each level of light intensity, tracking the angle of the shoot *θ*_tip_(*t*) = *θ*(*t, s* = *L*) over long periods of time (Fig. 2A, Supplementary Video S1), as described in the Methods. Fig. 2B shows average trajectories of tip angles *θ*_tip_ for a subset of light intensities (trajectories for all intensities, and an example of single trajectories in Fig. S2). After approximately 16 hours, plants reach a steady-state, and the average angle *θ*_f_ (highlighted in grey) represents the magnitude of the phototropic response, and clearly increases with light intensity.

**Fig. 2.**
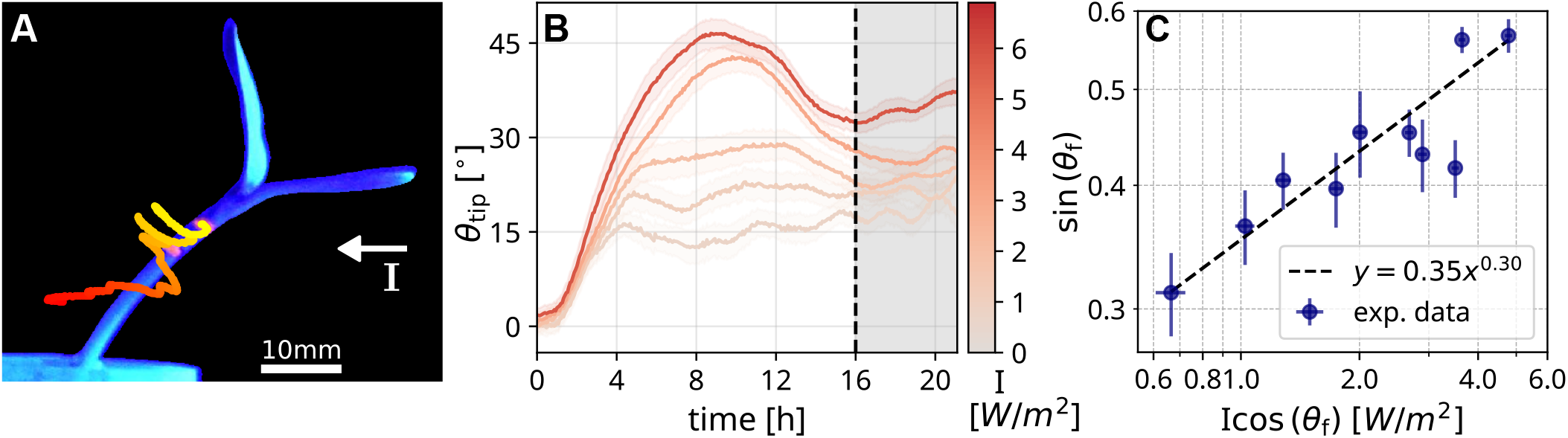
Determination of the phototropic transduction function. **(A) Quantification of phototropic bending**.Representative trajectories of fluorescent markers tracked over time during a phototropic response to a single light source (white arrow indicates light direction). The angle between two markers near the tip defines the time evolution of the tip angle *θ*_tip_(*t*) (video in Supplementary Video S1). **(B) Tip angle trajectories show distinct steady states for different light intensities**. Average tip angle trajectories *θ*_tip_(*t*) for increasing light intensities *I* (shaded regions around the average trajectories indicate standard error, 16 *< N <* 32; colorbar indicates light intensity). An example of individual trajectories is shown in Fig. S2A. After approximately 16 hours, seedlings reach a steady state that persists until the end of the experiment. We define *θ*_f_(*I*) as the average angle during this steady-state regime, which increases with light intensity. **(C) Determination of nonlinear transduction function**. Based on the planar steady-state relation between tip angle *θ*_f_ and light intensity *I* (Eq.5), we plot the measured output sin *θ*_f_ against the input variable *I* cos(*θ*_f_), which appears in the power-law relation. This allows direct extraction of the transduction function *ν*(*I*) (Eq.4) from the data. Fitting to the power-law form and substituting *T* = 0.1 yields *ν*_0_ */β* = 0.71 and a compressive exponent *α* = 0.30 (*R*^2^ = 0.73). Error estimation is detailed in SM A and Fig. S4.

Based on the phototropic dynamics for a single light source in Eq. 1, we find a 2D planar expression relating the steady-state angle of the tip *θ*_f_ to the transduced light and gravity signals (detailed in SM B), following sin 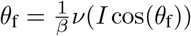. This provides an input-output relation for the transduction function, relating the measured output angle sin *θ*_f_ to the input signal *I* cos(*θ*_f_), the component of light intensity perpendicular to the organ. Substituting the power-law in Eq. 4 and assuming a transmission factor T yields:

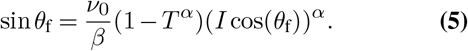

We note that if the angle between the directions of light and gravity is small, a similar vector derivation reproduces the equivalent *Resultant Angle* defined in (24) (detailed in SM D). We plot the measured output sin *θ*_f_ against the input signal *I* cos(*θ*_f_), and fit it to the power law following Eq. 5 (Fig. 2C). Substituting the measured value of the transmission factor *T* = 0.1 (see Methods) yields the prefactor *ν*_0_*/β* = 0.71, representing the ratio between phototropic and gravitropic sensitivities, and the power *α* = 0.30. A power *α <* 1 provides compressive action, enabling responses across a wide range of intensities, and has been measured for photosensors in a variety of organisms (51), as well as other sensory modalities which must cope with enormous ranges of energy.

### Bilateral lighting experiments corroborate vectorial summation model

Having established the functional form of the phototropic transduction function *ν*(*I*), we now proceed to experimentally corroborate the theoretical framework presented in Eqs. 1 and 3, describing the encoding and integration of spatial information regarding multiple light sources. We place sunflower seedlings between two parallel light sources, thus exposing them to two opposing light sources with intensities I_L_ and I_R_ from the left and right, respectively (Fig. 2A). Like before, the tropic responses are restricted to the plane defined by the two opposing light sources and the seedling, and we track the tip angles *θ*_tip_ over time. Fig. 3A shows the average tip trajectories *θ*_tip_(*t*) in the case of bilateral lighting, for a combination of light intensities in the range 0.7− 7.7 W*/*m^2^. We define the difference of intensities Δ*I* = *I*_R_− *I*_L_, with values in the range 0.25 −7.0 W*/*m^2^. As expected, we find that plants grow towards the direction with a higher light intensity, and the bigger the difference in intensity, the larger the steady-state angle *θ*_f_. However, this relation does not following a simple linear dependence on Δ*I*, as shown in Fig. 3B.

**Fig. 3.**
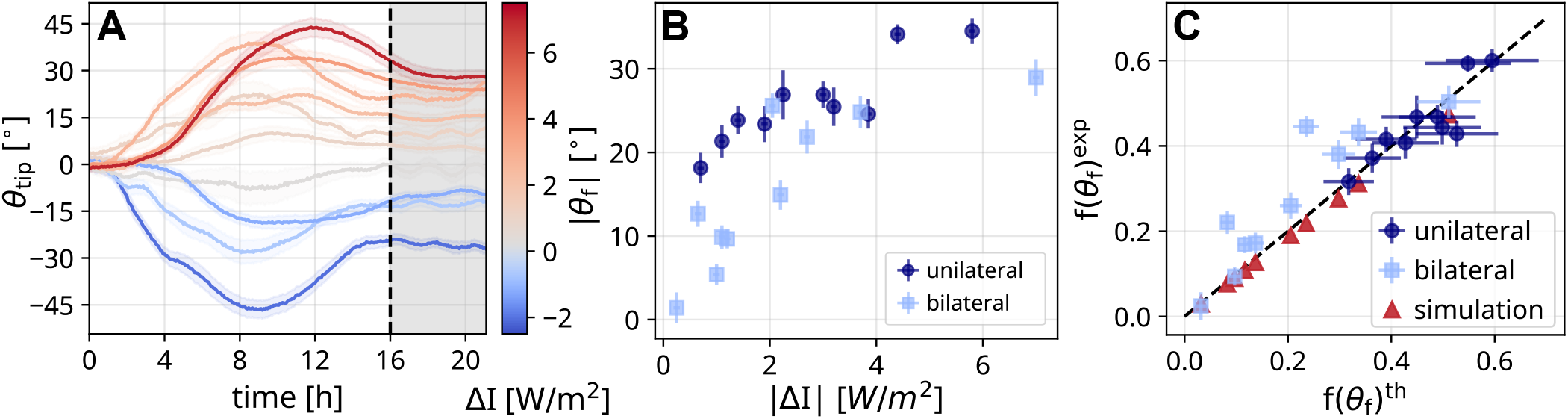
Experimental validation of vectorial integration under bilateral lighting. **(A) Bilateral lighting experiments**. Tip angle trajectories *θ*_tip_(*t*) for seedlings exposed to two opposing light sources from the left and right, with intensities *I*_L_ and *I*_R_ accordingly. The colorbar indicates the difference between intensities Δ*I* = *I*_L_ *I*_R_ (specific values in Methods). As in Fig. 2B, shaded regions around the average trajectories indicate standard error (19 *< N <* 36), and shaded grey area marks the steady state. *θ*_f_ increases with Δ*I*, and depends on its sign. **(B) Non-linear dependence of steady state angle on light difference**. Measured steady-state angles *θ*_f_ as a function of the light intensity difference |Δ*I*| for both unilateral and bilateral illumination. The response is monotonic but clearly nonlinear. Owing to the symmetry of the setup, absolute values are shown. **(C) Validation of steady-state relation for vectorial summation with bilateral lighting**. To test Eq.6, we compare its left- and right-hand sides. We define *f* (*θ*_f_) = sin *θ*_f_*/*(cos *θ*_f_)^*α*^, corresponding to the left-hand side of Eq.6. For each pair of light intensities *I*_L_, *I*_R_, we compute the theoretical value *f* ^th^(*θ*_f_) by substituting *I*_L_ and *I*_R_ into the right-hand side of Eq. 6. Model parameters *α* and *ν*_0_*/β* are taken from the fit in Fig.2C. We then plot *f* (*θ*_f_) against the experimentally measured value *f* ^exp^(*θ*_f_). Results are shown for unilateral (dark blue circles), bilateral (light blue squares), and simulated (orange triangles) conditions. Agreement along the identity line (dotted) indicates consistency between experiment and the model.

Similarly to the previous section, we tailor Eqs. 1 and 3 to describe the bilateral lighting setup, resulting in a 2D planar description (derivation detailed in the SM). In the specific case where light sources are opposite, I_L_ ∥ −I_R_, and in the same plane as the plant 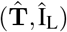, as described in Eq. 3, each side of the shoot is affected by both the direct and transmitted light. We therefore sum these, and define the effective light on each side. Substituting these light intensities in the relation for steady-state dynamics, as before, yields an expression relating the output steady-state angle *θ*_f_ to the magnitudes of the input, the two opposing light intensities *I*_L_ and *I*_R_:

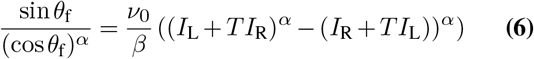

We note that substituting *I*_R_ = 0 reproduces the case for unilateral lighting in Eq. 5.

Next, we substitute the experimental values of *I*_L_ and *I*_R_, for both bilateral lighting and unilateral lighting experiments, yielding predictions for the steady-state angles. The predicted values *f* ^th^(*θ*_f_) ≡ sin(*θ*_f_)*/* cos(*θ*_f_)*α* are in good agreement with the experimentally measured values *f* ^ex^(*θ*_f_) (Fig. 3C). To further validate the model, we perform simulations using experimentally informed parameters under comparable bilateral lighting conditions, based on Eqs.1 and 3 (see Methods, Fig. S3). The simulations are consistent with both the theoretical predictions and experimental measurements, collectively providing quantitative support for the model of nonlinear vectorial representation and summation.

### Vectorial symmetry-breaking arguments predict plants do not maximize physical light

A key prediction of the vectorial model states that plants do not maximize light. A clear observation is that equal and opposite light sources effectively “cancel out”, so that the plant does not change its growth direction, as already shown in the bilateral lighting experiments for small values of Δ*I* (Fig. 3A). This observation is further articulated when a third weaker light source is added perpendicular to the plane defined by the two strong light sources. In this case the vectorial model predicts that since the two opposite vectors cancel out, the plant will grow towards the weaker perpendicular light which breaks the symmetry. We corroborate this experimentally (see inset of Fig. 4A), placing two opposite light sources with similar intensities (3.85 and 3.86 *±* 0.06 W*/*m^2^), and a third perpendicular light source with a weaker intensity (1.74 *±* 0.11 W*/*m^2^). Fig. 4A shows a polar distribution of average growth directions (see Methods). The average direction is 0.8° with a standard error 12.27°, clearly out of the plane defined by the two strong lights, towards the weaker light. This behavior is non-intuitive from a photosynthetic or energetic perspective, and establishes that plants do not maximize physical light.

**Fig. 4.**
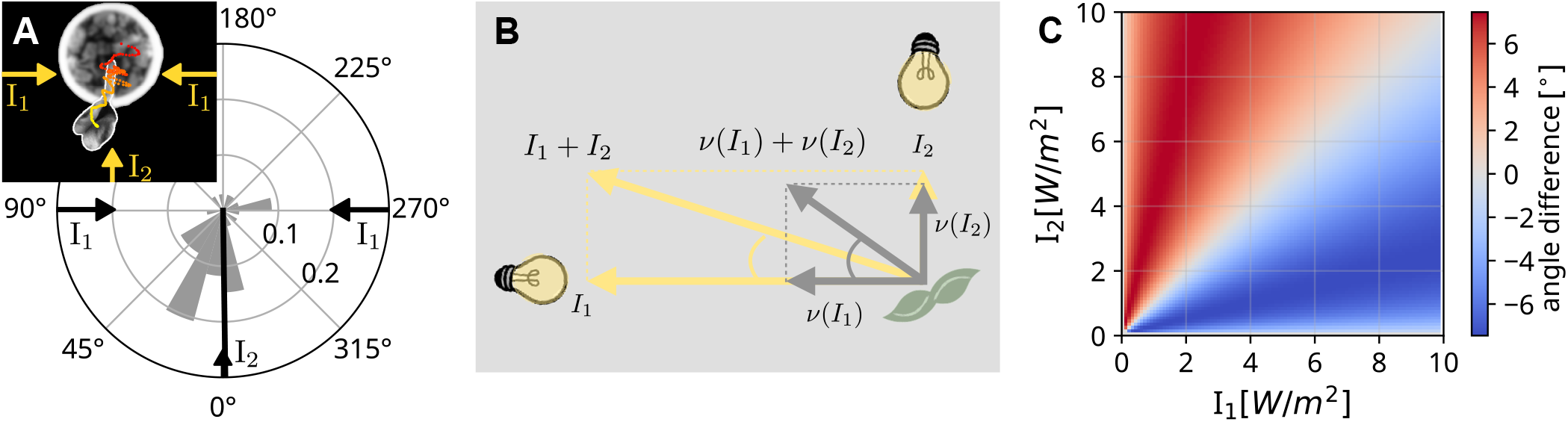
Vectorial integration and nonlinear distortions in multi-source illumination. **(A) Experimental demonstration of vectorial symmetry breaking under out-of-plane illumination**. Polar frequency distribution of final growth directions measured from a top view under out-of-plane illumination. Two equal and opposite light sources *I*_1_ (with similar values 3.86 *±* 0.06 W*/*m^2^ at −90^0^, and 3.85 *±* 0.06 W*/*m^2^ at +90^0^) are combined with a weaker perpendicular source *I*_2_ = 1.74 *±* 0.11 W*/*m^2^ at 0^0^. Despite the stronger opposing cues, seedlings grow toward the weaker symmetry-breaking source, demonstrating vector cancellation. The black line indicates the mean direction 0.8^°^ (standard error 12.26^°^). (inset) Experimental configuration and representative trajectory from a top-view recording. **(B) Conceptual illustration of distortion arising from nonlinear distributed sensing**. Two perpendicular light sources of unequal magnitude (yellow vectors). The physical total illumination is given by their vector sum. Because each signal is transduced nonlinearly and sensed locally across the surface, the vectorial sum of perceived or transducd signals (gray vectors) deviates in direction from the physical sum. **(C) Quantitative prediction of distortion between physical and perceived directions**. Angular difference between the direction of maximal physical illumination, obtained by integrating the incident light field over the shoot circumference (as defined in the main text), and the direction of maximal perceived illumination obtained from nonlinear distributed integration of transduced signals (Eq.7), computed for two perpendicular light sources as in (B). The colorbar indicates the angular deviation between these two directions. Deviations are symmetric in intensity ratio and vanish when one source is absent or when intensities are equal. In our parameter regime, maximal deviations are small (∼ 6^°^) and fall below experimental resolution (see Fig.S6 for additional examples).

### Non-linear perception in distributed sensory system leads to skewed responses

A direct consequence of the non-linear vectorial representation and integration of spatial information, validated here, is an inherent distortion of perceived illumination patterns. Namely, small differences arise between the perceived direction of total light illumination (integrated transduced signal), and the real, physical direction of maximal illumination. This can be graphically understood in Fig. 4B, a depiction of a top-view of a hypothetical experiment where a plant is exposed to two perpendicular light sources with magnitudes *I*_2_ *> I*_1_, represented vectorially by two yellow arrows. From the physical perspective the total light is equal to the vectorial sum of these two **I**_1_ + **I**_2_. However, since each signal is perceived separately, the magnitude of each signal is is transduced non-linearly (Eq. 4), altering the ratio between them (represented by grey arrows). Thus, the vectorial sum of the perceived stimuli, the direction of total light that the plant perceives, points in a different direction compared to the physical sum. This is effectively an optical illusion.

Since in this configuration the illuminations of the two light sources overlap on some parts of the shoot, a quantitative prediction of the distortion requires taking into account the geometry of photoreceptors distributed over the plant circumference. Light can no longer be approximated as a single effective vector; instead, it must be treated as a field acting locally on the surface, with its magnitude attenuated according to the local orientation of the surface, following Lambert’s cosine law (31, 52) (Fig. S5). The net incidence light is then 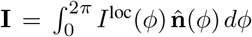, where *I*^loc^ (*ϕ*) is the light intensity reaching the angle *ϕ* along the cross-section (Eq. S.27, detailed derivation in SM C), and 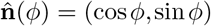 is the normal. The resultant direction of the illumination pattern over the circumference is then Î = **I***/* ∥**I**∥ . The net perceived signal is

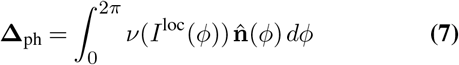

and the resultant direction of the perceived illumination pattern is then 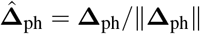 . This is the general scalar field version of Eq. 3.

In Fig. 4C we calculate the difference between the direction of physical net incidence light **Î**, and the perceived direction 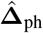, for two perpendicular light sources *I*_1_ and *I*_2_ (Fig. 4B), across a range of intensities. Deviations are symmetric in *I*_1_ and *I*_2_, and vanish when *I*_1_ = 0, *I*_2_ = 0, or *I*_1_ = *I*_2_, where both vectors point at 45°. Fig. S6 shows the full directional dependence for selected values of *I*_2_. The predicted angular differences are small, with maximal values of ∼6° for different intensity ratios. These values are below the resolution of our experiments, as they are smaller than the intrinsic variability in measured growth direction. In particular, the standard error of the growth direction (Fig.4A) is 12.27°, and the distribution of standard deviation of the angles during the last three hours (Fig.S7) has a median of 16.64°. This variability arises primarily from inherent plant movements termed circumnutations (see representative trajectories in Fig. S8), and therefore sets a practical detection limit.

## Discussion

We developed an experimentally informed framework describing how plants integrate multiple spatially distributed light cues. Our results show that phototropic responses are governed not by the physical vectorial sum of incident light, but by the vectorial sum of locally transduced signals distributed across the shoot circumference. Because sensory transduction is nonlinear and light is sensed over an extended curved surface, this integration rule differs systematically from physical summation.

One consequence of this integration rule, which we validated experimentally, is that symmetric opposite high-intensity light sources cancel at the level of transduced signals, allowing a weaker symmetry-breaking cue to dominate the response. We confirmed this prediction by introducing a third out-of-plane light source, demonstrating that seedlings grow toward the weaker cue despite the presence of stronger opposing stimuli. This behavior establishes that plants do not maximize physical light intensity, but instead follow the vectorially integrated representation of locally encoded signals. A second consequence follows directly from the same geometric formulation. Nonlinear local encoding combined with circumferential sensing predicts that ∑_*i*_ *ν*(*I*_*i*_) ≠ *ν*(∑_*i*_ *I*_*i*_); in other words, there are systematic discrepancies between the direction of maximal physical illumination and the direction of growth. In our parameter regime these predicted distortions are small and fall below experimental detection due to intrinsic variability such as circumnutation. Nevertheless, they arise generically from distributed nonlinear sensing and constitute an effective optical illusion at the level of internal representation. In spite of these distortions, tropic sensing in natural environments is likely optimized for robust directional resolution under heterogeneous illumination rather than for maximizing instantaneous photon capture. Compressive nonlinear transduction facilitates responses across a wide range of intensities, from shaded canopies to direct sunlight. Furthermore, we note that in plants the vectorial representation of stimuli, and their integration, can be naturally encoded in spatial gradients of growth rate across the organ cross-section (Fig.1A). In our framework, each directional stimulus induces a differential growth pattern corresponding to a gradient in axial growth rate, which can be interpreted as a local vector encoded at the tissue level. Because such growth-rate gradients add linearly, their superposition implements the vectorial sum of transduced signals (detailed mathematically in SMD), providing a physical realization of distributed cue integration.

Mathematical models of multi-cue integration in systems such as chemotactic cells have successfully described competing gradients through reaction–diffusion and LEGI-type frameworks (12, 13), as well as nonlinear receptor-level integration in bacteria (3, 53). These models capture biased and hierarchical responses through intracellular signaling dynamics and adaptation. In contrast, the mechanism identified here emerges from the geometry of distributed sensing and nonlinear local transduction, independent of specific biochemical circuitry. In our minimal model, systematic distortions between physical stimulus fields and integrated directional responses arise as a structural consequence of surface-based integration rather than molecular signaling.

Plants provide a particularly transparent example of a multicellular system in which sensing and integration are fully decentralized. Because the mechanism identified here relies only on distributed sensing over extended geometries and nonlinear local encoding, it is not plant-specific. Similar geometric effects may operate in other biological systems that integrate spatially resolved cues across extended surfaces, including chemotactic cells and other decentralized organisms. In these systems, external chemical or mechanical gradients are sampled across a cell membrane or tissue surface and converted into a polarized internal state, such as asymmetric cytoskeletal activity or traction forces, that defines a global migration or growth direction. More broadly, the geometric framework developed here may inform engineered systems that employ distributed sensing without centralized control, including soft robotic and embodied sensing platforms.

## Materials and Methods

### Plant material and growth conditions

Seedlings of *Helianthus annus* from the “HAEMEK-6” verity were grown in 50 mL tubes filled with vermiculite in a growth chamber maintained at 25 ^°^C with a 12 h light–12 h dark cycle. Seedlings were left to grow for approximately one week at which point plants with a uniform size of 1-3 cm were selected and transferred into the experiment room. Fluorescent color markers were painted on the plants in order to simplify the tracking process. Finally the plants were placed in the setup resting in darkness for one hour before the beginning of the experiment, which in turn lasted at least 21 h. To avoid light signaling, the selection and marking process was done at the end of the dark period of the cycle and under dim greenlight conditions.

### Transmittance measurement

To assess light transmission through the plant, we designed and 3D-printed a custom sleeve for our Ophir light sensor (Fig. S9). The sleeve allowed for precise placement of a seedling stem to occlude the light. Light intensity was then recorded under both obstructed and unobstructed conditions to calculate the percentage of transmitted light. This process was repeated across multiple distances and intensities of light sources, using LED light sources as described above. A simple average of all measurements was calculated and used, ultimately producing a transmission factor of *T* = 9.8% *±*0.8% (mean *±*standard error, *N* = 31).

### Experimental setup

In order to quantify plants’ responses to different light signals, we built an experimental setup for unilateral and bilateral lighting, simultaneously exposing five plant shoots to one light source, or two opposite light sources with different intensities. We tracked the phototropic movement throughout, using the image analysis procedure described below. The setup consists of two custom-built LED panels, using blue LED strips with a wavelength of 450nm. Panels were placed facing each other on two sides of the plant holder grid. The light intensity was measured before the beginning of the experiment using a c-50 Ophir-Optronics power meter at each of the plant locations. In bilateral lighting experiments, specific intensity pairs were as follows (*I*_L_, *I*_R_): (7.7,0.7), (4.7,1.0), (3.7, 1.5), (0.95, 3), (2.9,1.9), (1.3, 1.95), (1.75, 1.5), (2.25, 1.15), (1.8, 3.0), (3.5, 0.8) W*/*m^2^ with corresponding |Δ*I*| : 7.0, 3.7, 2.2, 2.05, 1.0, 0.65, 0.25, 1.1, 1.2, 2.7 W*/*m^2^. The schematics of the experimental setup can be found in Fig. S1. For the 3D experiments, we introduced a third LED panel positioned orthogonal to the plane created by the first two light sources. Here we used two equal light intensities on opposite sides of the plant *I*_1_ = 3.86 *±*0.06 W*/*m^2^, *I*_2_ = 3.85 *±*0.06 W*/*m^2^, and a third weaker light source *I*_3_ = 1.74 *±*0.11 W*/*m^2^ perpendicular to the others. The plant’s movement was then recorded from above during 21 h, using the image analysis procedure described below (N=34).

### Image and data analysis

For unilateral and bilateral experiments, images were taken from a side view. To quantify phototropic movement on the basis of the recorded images, we extracted the location of two points painted at the tip of the plant and a few millimeter below (Supplementary Video S1); to this end, we used basic image analysis and a Python implementation of the CSRT tracker (54) from the open source OpenCV library (55). These points were then used to calculate the angle at the tip of the plant *θ*(*t*). The steady state angle *θ*_f_ is the average angle during the steady state regime. For out-of-plane three-point illumination experiments, images were acquired from a top view (example in inset of Fig.4A). Because painted markers were not visible in this configuration, we tracked the leaves instead: a YOLOv8 model (56), trained on manually labeled images, was used to segment the leaf region in each frame of the time-lapse. A single (x,y) position was then extracted for each frame as the center of mass of the segmented region. The base of the plant was manually annotated. The growth direction was defined as the angle between the base and the mean (x,y) position over the final 3 h of the experiment. Representative trajectories are shown in Fig.S8.

### Simulations

Simulations were performed using the code published in (25), which includes explicit axial growth, modified by new phototropic terms that follow Eq. 4 and the transmission factor *T* = 0.1. The geometry and growth parameters of the simulated shoots were estimated: *R* = 1.25 mm was measured using calipers for the radius of the cross section. *L*_0_ = 20 mm was estimated for the initial length, and an initial straight condition. Axial growth was assumed to be uniform within a finite growth zone of length *L*_*gz*_ = 20 mm, with constant relative growth rate 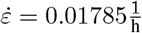. The tropic dynamics was then integrated over ∼23 hours, so that the final length is *L*_*f*_ = 28.2 mm. The value of the ratio between phototropic and gravitropic sensitivities *ν*_0_*/β* = 0.713 was taken from the experiments reported herein (Fig. 2C), and *α* = 0.297. Specific values for gravitropic sensitivity are *β* = 2, and autotropic sensitivity *γ* = 5.

## ACKNOWLEDGEMENTS

We thank Amir Ohad, Agueda de la Vega, and Alessia Perilli for helpful conversations. We thank Karen Marron for scientific editing of the paper.

## AUTHOR CONTRIBUTIONS

A.K. and M.R.: experimental design; A.K. methodology, carried out all experiments, experimental data curation; A.P: developed simulation framework, carried out simulations, simulation data curation; A.K., A.P and Y.M: formal analysis, conceptualization and investigation, visualization, writing original draft; A.K., M.R., A.P and Y.M: review, and editing, Y.M. funding acquisition, supervision, and resources.

## DATA AVAILABILITY

Datasets and analysis scripts used to generate trajectories are openly available via Zenodo: https://zenodo.org/uploads/19110267 (57). Includes spreadsheets of experimental data extracted from images, Python scripts for running simulations, data analysis, and reproduction of all figures presented in the article.

