## SUPPLEMENTARY MATERIAL for "Nonlinear distributed sensing of light patterns leads to perceptual distortions in plants"

### Supplementary Materials for
### Nonlinear distributed sensing of light patterns leads to perceptual distortions in plants

**Ahron Kempinski, Amir Porat, Mathieu Rivière, Yasmine Meroz***

#### This PDF file includes:

Supplementary Text
Figures S1 to S9
Captions for Movies S1

#### Other Supplementary Materials for this manuscript:

Movies S1

**A. Error estimation for Fig. 2C and Fig. 3C.** For Fig. 2C we calculate the errors for the fitting of unilateral light experiments using Eq. 5 and Eq. 4 as follows:

$$\sin(\theta_{\rm f}) = I\cos(\theta_{\rm f}) \tag{S.1}$$

$$y \equiv \sin(\theta_{\rm f});\;\; x \equiv I\cos(\theta_{\rm f}) \tag{S.2}$$

$$dx = \sqrt{\left(\frac{\partial x}{\partial I}\right)^2 (dI)^2 + \left(\frac{\partial x}{\partial \theta_{\rm f}}\right)^2 (d\theta_{\rm f})^2} \tag{S.3}$$

$$dx = \sqrt{(\cos^2(\theta_{\rm f})\,(dI)^2 + (-I\sin(\theta_{\rm f}))^2 (d\theta_{\rm f})^2} \tag{S.4}$$

$$dy = \sqrt{\left(\frac{\partial y}{\partial \theta_{\rm f}}\right)^2 (d\theta_{\rm f})^2} \tag{S.5}$$

$$dy = \sqrt{\cos^2(\theta_{\rm f})\,(d\theta_{\rm f})^2} = \cos(\theta_{\rm f})\,(d\theta_{\rm f}) \tag{S.6}$$

For Fig. 3C we use Eq. 6 and change it to represent experimental values and theoretical predictions:

$$\frac{\sin(\theta_{\rm f})}{\cos(\theta_{\rm f})^b} = \frac{\nu_0}{\beta}\left((I_{\rm L} + T * I_{\rm R})^b - (I_{\rm R} + T * I_{\rm L})^b\right) \tag{S.7}$$

$$y \equiv \frac{\sin(\theta_{\rm f})}{\cos(\theta_{\rm f})^b};\;\; x \equiv \frac{\nu_0}{\beta}\left((I_{\rm L} + T * I_{\rm R})^b - (I_{\rm R} + T * I_{\rm L})^b\right) \tag{S.8}$$

$$dx = \sqrt{\left(\frac{\partial x}{\partial I_{\rm L}}\right)^2 (dI_{\rm L})^2 + \left(\frac{\partial x}{\partial I_{\rm R}}\right)^2 (dI_{\rm R})^2 + \left(\frac{\partial x}{\partial T}\right)^2 (dT)^2} \tag{S.9}$$

$$\frac{\partial x}{\partial I_{\rm L}} = \frac{\nu_0}{\beta}\left(b(I_{\rm L} + TI_{\rm R})^{b-1} - b(I_{\rm R} + TI_{\rm L})^{b-1}T\right) \tag{S.10}$$

$$\frac{\partial x}{\partial I_{\rm R}} = \frac{\nu_0}{\beta}\left(b(I_{\rm L} + TI_{\rm R})^{b-1}T - b(I_{\rm R} + TI_{\rm L})^{b-1}\right) \tag{S.11}$$

$$\frac{\partial x}{\partial T} = \frac{\nu_0}{\beta}\left(b(I_{\rm L} + TI_{\rm R})^{b-1}I_{\rm R} - b(I_{\rm R} + TI_{\rm L})^{b-1}I_{\rm L}\right) \tag{S.12}$$

$$dy = \sqrt{\left(\frac{\partial y}{\partial \theta_{\rm f}}\right)^2 (d\theta_{\rm f})^2} \tag{S.13}$$

$$\frac{\partial y}{\partial \theta_{\rm f}} = (\cos(\theta_{\rm f}))^{-b-1}\left(b\sin^2(\theta_{\rm f}) + \cos^2(\theta_{\rm f})\right) \tag{S.14}$$

**B. Relation for steady state tip angle.** We define the direction of gravity as $\hat{\boldsymbol{g}} = -\hat{\mathbf{y}}$, and the light source $\mathbf{I} = -I\hat{\mathbf{x}}$ (Fig. 1A). Assuming the initial condition of the shoot is straight, parallel to the $\hat{\mathbf{y}}$ direction, the tropic dynamics are restricted to the $\hat{\mathbf{x}} - \hat{\mathbf{y}}$ plane, allowing for a simplified formulation.
The component of the signal $\mathbf{I}$ perpendicular to the organ direction $\hat{\mathbf{T}}$ follows

$$\mathbf{I}^{\perp} = \hat{\mathbf{T}} \times (\mathbf{I} \times \hat{\mathbf{T}}). \tag{S.15}$$

Substituting $\mathbf{I} = -I\hat{\mathbf{x}}$, and following geometrical arguments, leads to an expression for the magnitude:

$$I^{\perp} = |\mathbf{I}^{\perp}| = I\sin(\frac{\pi}{2} - \theta) = I\cos(\theta), \tag{S.16}$$

where $\frac{\pi}{2} - \theta(s,t)$ is the local angle between the tangent $\hat{\mathbf{T}}$ and the signal direction $\hat{\mathbf{I}}$, and $\theta(s,t)$ is the local angle of the organ from the vertical (Fig. 1A).

The unit vector of the signal direction is then:

$$\hat{\mathbf{I}}^{\perp} = \frac{\mathbf{I}^{\perp}}{I^{\perp}} = \frac{\hat{\mathbf{T}} \times (-I\hat{\mathbf{x}} \times \hat{\mathbf{T}})}{I\cos(\theta)} = \frac{1}{\cos(\theta)}\hat{\mathbf{T}} \times (-\hat{\mathbf{x}} \times \hat{\mathbf{T}}), \tag{S.17}$$

Substituting Eqs. S.16-S.17 into Eq. 2 yields:

$$\boldsymbol{\Delta}_{\text{ph}} = -\nu(|\mathbf{I}^{\perp}|)\hat{\mathbf{I}}^{\perp} = \frac{\nu(I\cos(\theta))}{\cos(\theta)}\hat{\mathbf{T}} \times (\hat{\mathbf{x}} \times \hat{\mathbf{T}}) \tag{S.18}$$

The differential growth associated with gravitropism, responding to $\mathbf{g} = -g\hat{\mathbf{y}}$ follows:

$$\boldsymbol{\Delta}_{\text{gr}} = -\beta\hat{\mathbf{g}}^{\perp} = \beta\hat{\mathbf{T}} \times (\hat{\mathbf{y}} \times \hat{\mathbf{T}}) \tag{S.19}$$

For simplicity, we focus on the tip, $s = L$, and assume that it is straight with $\kappa(L) = 0$, such that autotropism is negligible with $\boldsymbol{\Delta}_{\text{au}}(L) = 0$.
Substituting the expressions Eqs. S.18 and S.19 into the expression for differential growth $\boldsymbol{\Delta} = \boldsymbol{\Delta}_{\text{ph}} + \boldsymbol{\Delta}_{\text{gr}} + \boldsymbol{\Delta}_{\text{au}}$ (as in Eq. 1), and applying the identity for cross products $\mathbf{A} \times (\mathbf{B} \times \mathbf{C}) = (\mathbf{A} \cdot \mathbf{C})\mathbf{B} - (\mathbf{A} \cdot \mathbf{B})\mathbf{C}$, finally yields:

$$\boldsymbol{\Delta} = \frac{\nu(I\cos(\theta))}{\cos(\theta)}(\mathbf{x} - (\mathbf{T} \cdot \mathbf{x})\mathbf{T}) + \beta(\mathbf{y} - (\mathbf{T} \cdot \mathbf{y})\mathbf{T}), \tag{S.20}$$

Projecting Eq.S.20 on $\hat{\mathbf{x}}$ or $\hat{\mathbf{y}}$, and recalling that $\mathbf{T} \cdot \mathbf{x} = \sin\theta$ and $\mathbf{T} \cdot \mathbf{y} = \cos\theta$ (Fig. 1A) yields:

$$\begin{aligned} \boldsymbol{\Delta} \cdot \hat{\mathbf{x}} &= \frac{\nu(I\cos(\theta))}{\cos(\theta)}(\mathbf{x} - (\mathbf{T} \cdot \mathbf{x})\mathbf{T}) \cdot \hat{\mathbf{x}} + \beta(\mathbf{y} - (\mathbf{T} \cdot \mathbf{y})\mathbf{T}) \cdot \hat{\mathbf{x}} = \\ &= \frac{\nu(I\cos(\theta))}{\cos(\theta)}(1 - (\mathbf{T} \cdot \mathbf{x})^2) + \beta(0 - (\mathbf{T} \cdot \mathbf{y})(\mathbf{T} \cdot \mathbf{x})) = \\ &= \frac{\nu(I\cos(\theta))}{\cos(\theta)}\cos^2(\theta) - \beta\cos(\theta)\sin(\theta) \end{aligned} \tag{S.21}$$

In the case of the steady-state solution, the organ has reached its final orientation described by the final tip angle $\theta_{\text{f}}$. The curvature remains constant in time, and thus Eq. 1 goes to zero, such that $\boldsymbol{\Delta} = 0$, i.e. there is no differential growth. Equating Eq. S.21 to zero finally yields the expression for the tip angle at steady state:

$$\boxed{\sin\theta_{\text{f}} = \frac{\nu(I\cos(\theta_{\text{f}}))}{\beta}} \tag{S.22}$$

Next, following the experimental setup, we consider two light sources directed at the plant shoot from opposite sides, such that the tropic response is again restricted to the $\hat{\mathbf{x}} - \hat{\mathbf{y}}$ plane. We define the light sources $\mathbf{I}_{\text{L}} = I_{\text{L}}\hat{\mathbf{x}}$ and $\mathbf{I}_{\text{R}} = -I_{\text{R}}\hat{\mathbf{x}}$ arbitrarily as coming from the left and right, respectively.
Following Eqs. S.16 and S.17, the components of the light signal perpendicular to the organ follow:

$$\mathbf{I}_L^{\perp} = I_L^{\perp}\hat{\mathbf{I}}_L^{\perp} = I_{\text{L}}\cos(\theta)\left(\frac{1}{\cos(\theta)}\hat{\mathbf{T}} \times (\hat{\mathbf{x}} \times \hat{\mathbf{T}})\right), \tag{S.23}$$

$$\mathbf{I}_R^{\perp} = I_R^{\perp}\hat{\mathbf{I}}_R^{\perp} = I_{\text{R}}\cos(\theta)\left(\frac{1}{\cos(\theta)}\hat{\mathbf{T}} \times (-\hat{\mathbf{x}} \times \hat{\mathbf{T}})\right), \tag{S.24}$$

Substituting Eqs. S.23,S.24 into Eq. 2 yields:

$$\begin{aligned} \boldsymbol{\Delta}_{\text{ph}} &= -\sum_{\text{i}}\nu(|\mathbf{I}_{\text{i}}^{\perp}|)\hat{\mathbf{I}}_{\text{i}}^{\perp} = -\nu(I_{\text{L}}^{\perp})\hat{\mathbf{I}}_L^{\perp} - \nu(I_{\text{R}}^{\perp})\hat{\mathbf{I}}_R^{\perp} = \\ &= (-\nu(I_{\text{L}}\cos\theta) + \nu(I_{\text{R}}\cos\theta))\frac{\hat{\mathbf{T}} \times (\hat{\mathbf{x}} \times \hat{\mathbf{T}})}{\cos(\theta)} \end{aligned} \tag{S.25}$$

Following the calculation for a single light signal in Eq. S.21, the relation between the steady state tip angle $\theta_{\text{f}}$ to the light and gravity signals is:

$$\boxed{\sin\theta_{\text{f}} = \frac{\nu(I_{\text{L}}\cos\theta_{\text{f}}) - \nu(I_{\text{R}}\cos\theta_{\text{f}})}{\beta}} \tag{S.26}$$

substituting: $I_{\text{L}} \to I_{\text{L}} + TI_{\text{R}}$ and $I_{\text{R}} \to I_{\text{R}} + TI_{\text{L}}$ to account for light transmission effects then gives Eq. 6 in the main text. In a similar fashion, substituting $I_{\text{L}} \to I$ and $I_{\text{R}} \to TI$ gives Eq. 5.

**C. Angular attenuation.** Since in this configuration the illumination of both light sources overlap on some parts of the shoot, a quantitative prediction of the distortion requires to take into account the geometry of photoreceptors distributed over the plant circumference. Light can no longer be approximated as a single effective vector; instead, it must be treated as a field acting locally on the surface, with its magnitude attenuated according to the local orientation of the surface.

Let us assume the cross-section of an organ is described with a polar angle $\phi$, and that the direction of a light stimulus of magnitude $I_{\mathrm{i}}$ is defined by the angle $\phi_{\mathrm{i}}$. For a cylindrical surface, the local light intensity on the *front* half of the cylinder, follows Lambert's cosine law, decreasing as the cosine of the angle between the incident light direction and the local surface normal (31, 52) (Fig. S5). The *back* half perceives a symmetric profile of the transmitted light. Thus, the perceived light intensity at an angle $\phi$ along the cross-section follows:

$$I^{\mathrm{loc}}(\phi) = \sum_{\mathrm{i}} \Big[ I_{\mathrm{i}} \max\{0, \cos(\Delta\phi_{\mathrm{i}})\} + T I_{\mathrm{i}} \max\{0, -\cos(\Delta\phi_{\mathrm{i}})\} \Big] \tag{S.27}$$

where $\Delta\phi_{\mathrm{i}} = \phi - \phi_{\mathrm{i}}$ is the angle relative to the stimulus angle.

For clarity, we break this calculation down first in the case of a single light source. Let us consider the half of the shoot circumference facing the light source, $-\frac{\pi}{2} < \phi - \phi_{\mathrm{s}} < \frac{\pi}{2}$. We recall that light is transduced locally. We define the vector of the transduced signal at each point along the circumference $\vec{\nu}(\phi)$, following:

$$\vec{\nu}(\phi) = \nu(I\cos(\phi - \phi_{\mathrm{s}})) \cdot (\cos(\phi), \sin(\phi)), \quad -\frac{\pi}{2} < \phi - \phi_{\mathrm{s}} < \frac{\pi}{2} \tag{S.28}$$

For each point the light comes into contact with the shoot, there is a transmitted signal on the opposite side:

$$\vec{\nu}(\phi) = (\nu(I\cos(\phi - \phi_{\mathrm{s}})) - \nu(TI\cos(\phi - \phi_{\mathrm{s}})) \cdot (\cos\phi, \sin\phi), \quad -\frac{\pi}{2} < \phi - \phi_{\mathrm{s}} < \frac{\pi}{2} \tag{S.29}$$

We now sum the total transduced signal, calculating the total x and y components:

$$\begin{aligned} \nu_x^{\mathrm{tot}} &= \int_{-\pi/2}^{\pi/2} \nu_x(\alpha)\cos(\alpha + \phi_{\mathrm{s}})d\alpha \\ &= \int_{-\pi/2}^{\pi/2} (\nu(I\cos\alpha) - \nu(TI\cos\alpha))\cos(\alpha + \phi_{\mathrm{s}})d\alpha \end{aligned} \tag{S.30}$$

$$\nu_y^{\mathrm{tot}} = \int_{-\pi/2}^{\pi/2} (\nu(I\cos\alpha) - \nu(TI\cos\alpha))\sin(\alpha + \phi_{\mathrm{s}})d\alpha \tag{S.31}$$

In the case of multiple light sources, we write:

$$\nu_x^{\mathrm{tot}}\hat{\mathbf{x}} + \nu_y^{\mathrm{tot}}\hat{\mathbf{y}} = \int_0^{2\pi} \nu\left(\sum_i I_i\left(f_i(\theta)H(f_i(\theta)) + Tf_i(\theta)H(-f_i(\theta))\right)\right)\hat{\mathbf{n}}(\theta)d\theta \tag{S.32}$$

where $i$ sums over different sources, $\hat{\mathbf{n}}$ is the outfacing normal to the epidermis of each cross section, $f_i(\theta) = \hat{\mathbf{n}}(\theta) \cdot (-\hat{\mathbf{I}}_i^{\perp})$, and $H$ is the Heaviside function. Since $\nu$ is a non linear function, the sum of sources from general different directions cannot be represented in vector form as in Eq. 2. Eq. 2 therefore serves as an approximation and is exact in the case of one source or two antiparallel sources.

**D. Small angle approximation reproduces steady-state angle for single light source.** We note that if the angle between directions of light and gravity is small, our vector formalism reproduces the *Resultant Angle* defined in (24) as the angle that balances two tropic factors. To show this, we assume gravitropism and phototropism such that

$$\boldsymbol{\Delta} = \boldsymbol{\Delta}_{\mathrm{gr}} + \boldsymbol{\Delta}_{\mathrm{ph}} = \nu\hat{\mathbf{T}} \times (\hat{\mathbf{n}} \times \hat{\mathbf{T}}) + \beta\hat{\mathbf{T}} \times (\hat{\mathbf{y}} \times \hat{\mathbf{T}}) \tag{S.33}$$

where the direction of light is defined using an angle $\alpha$ such that $\hat{\mathbf{n}} = \sin(\alpha)\hat{\mathbf{x}} + \cos(\alpha)\hat{\mathbf{y}}$. This gives:

$$\boldsymbol{\Delta} = \nu\sin(\alpha)\hat{\mathbf{T}} \times (\hat{\mathbf{x}} \times \hat{\mathbf{T}}) + (\nu\cos(\alpha) + \beta)\hat{\mathbf{T}} \times (\hat{\mathbf{y}} \times \hat{\mathbf{T}}) \tag{S.34}$$

Assuming steady state with $\mathbf{\Delta}\cdot\hat{\mathbf{x}}=0$ at the tip gives:

$$0=\mathbf{\Delta}\cdot\hat{\mathbf{x}}=\nu\sin(\alpha)\cos^2(\theta_\mathrm{f})-(\nu\cos(\alpha)+\beta)\sin(\theta_\mathrm{f})\cos(\theta_\mathrm{f}) \tag{S.35}$$

which reproduces the expression found in (24):

$$\theta_\mathrm{f}\approx\tan(\theta_\mathrm{f})=\frac{\nu\sin(\alpha)}{\beta+\nu\cos(\alpha)}\approx\frac{\nu}{\beta+\nu}\alpha \tag{S.36}$$

where the first and last approximations are small angle approximations.

**Mathematical derivation showing plants can encode non-linear transformations of vectors, using differential growth gradients.** Here we describe a more general version of Eq. 1 with an explicit account for growth (e.g. in (25, 58)). Using the same definitions as in the main text, we introduce growth, denoting $\dot{\varepsilon}(\boldsymbol{\rho})$ the local axial relative growth rate at a point $\boldsymbol{\rho}=\rho(\cos(\phi),\sin(\phi))$ in the cross section, in polar coordinates. We then define the growth rate of the centerline $\dot{\varepsilon}(0)\equiv\dot{\varepsilon}_0$ as the average across the circumference of the circular cross section with radius $R$: $\dot{\varepsilon}_0=(\dot{\varepsilon}(\boldsymbol{R})+\dot{\varepsilon}(-\boldsymbol{R}))/2$. For small curvatures ($|R\boldsymbol{\kappa}|\ll 1$) this guarantees that growth is compatible and the cross section remains planar. The tropic dynamics of plant organs in response to external stimuli is driven by a differential growth rate across the plant cross section, which we define as a vector $\mathbf{\Delta}$ describing the difference across the organ circumference, such that (25, 58):

$$\mathbf{\Delta}\cdot\hat{\boldsymbol{\rho}}=\frac{\dot{\varepsilon}(-R\hat{\boldsymbol{\rho}})-\dot{\varepsilon}(R\hat{\boldsymbol{\rho}})}{\dot{\varepsilon}(-R\hat{\boldsymbol{\rho}})+\dot{\varepsilon}(R\hat{\boldsymbol{\rho}})}\approx-\frac{R}{\dot{\varepsilon}_0}\boldsymbol{\nabla}\dot{\varepsilon}\cdot\hat{\boldsymbol{\rho}} \tag{S.37}$$

where $\hat{\boldsymbol{\rho}}$ is a general direction on the cross section plane, and in the last approximation we use the average growth rate and its gradient on the centerline.

The resulting growth-driven dynamics are then described by relating the differential growth vector to the co-rotational material time derivative of the curvature vector in the plane of the organ cross-section:

$$\frac{D\boldsymbol{\kappa}}{Dt}=\frac{\dot{\varepsilon}_0}{R}\hat{\mathbf{T}}\times\mathbf{\Delta}. \tag{S.38}$$

Here, the material time derivative follows $\frac{D}{Dt}=\frac{\partial s}{\partial t}\frac{\partial}{\partial s}+\frac{\partial}{\partial t}$, where $\frac{\partial s}{\partial t}=v(s,t)=\int^s\dot{\varepsilon}(u,t)du$ is the local velocity due to growth. For simplicity, we omit the explicit dependence on position s along the centerline and time t.

In tropisms, the differential growth vector $\mathbf{\Delta}$ is dictated by external directional stimuli, such as light and gravity.

Concretely, the differential growth of a plant exposed to a single light source and gravity, includes a sum of the respective contributions $\mathbf{\Delta}_{\mathrm{ph}}$ and $\mathbf{\Delta}_{\mathrm{gr}}$, and $\mathbf{\Delta}_{\mathrm{au}}$ autotropism or proprioception, as described in detail in the main text, following:

$$\mathbf{\Delta}=-\nu(I^\perp)\hat{\mathbf{I}}^\perp-\beta\hat{\mathbf{g}}^\perp-\gamma R\kappa\hat{\mathbf{N}} \tag{S.39}$$

where for a general signal $\mathbf{u}$ this component is defined as $\mathbf{u}^\perp=\mathbf{u}-\hat{\mathbf{T}}(\mathbf{u}\cdot\hat{\mathbf{T}})=\hat{\mathbf{T}}\times(\mathbf{u}\times\hat{\mathbf{T}})$. Together, Eqs S.38 and S.39 are the equivalent of Eq. 1 with an explicit account for growth, which in many cases can be neglected (58).

The additive decomposition of the differential growth vectors can be justified by assuming that the local axial growth rate $\dot{\varepsilon}$ is a function of various local properties $\{\mu_i\}$, such that $\dot{\varepsilon}(\mu_1,...,\mu_N)$, where each property $\mu_i$ is allowed to vary spatially in the organ. The properties $\{\mu_i\}$ may symbolize any field that can affect the local growth rate, such as distributions of various morphogens, proteins, or other physical properties such as mechanical stress, Turgor pressure or cell wall properties. Following Eq. S.37 and the chain rule, these dependencies give:

$$\mathbf{\Delta}=-\frac{R}{\dot{\varepsilon}_0}\boldsymbol{\nabla}\dot{\varepsilon}(\mu_1,...\mu_N)=-\frac{R}{\dot{\varepsilon}_0}\sum_{i=1}^{N}\frac{\partial\dot{\varepsilon}}{\partial\mu_i}\boldsymbol{\nabla}\mu_i\equiv\sum_i\mathbf{\Delta}_i \tag{S.40}$$

Therefore, the additive decomposition of the differential growth vector can be related to a multi variable growth law.

This provides a basis for understanding how plants can encode non-linear transformations of vectors, and sum these vectorially.

#### E. Supplementary figures.

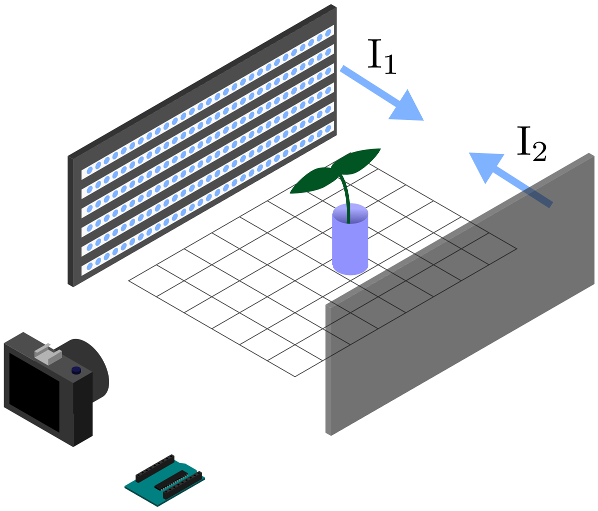


**Fig. S1. Schematic of the experimental setup.** Plants were placed between two two adjustable light sources each providing a distinct light intensity from opposing sides. Light output was controlled via an Arduino micro-controller, while the plants response was recorded using a camera.

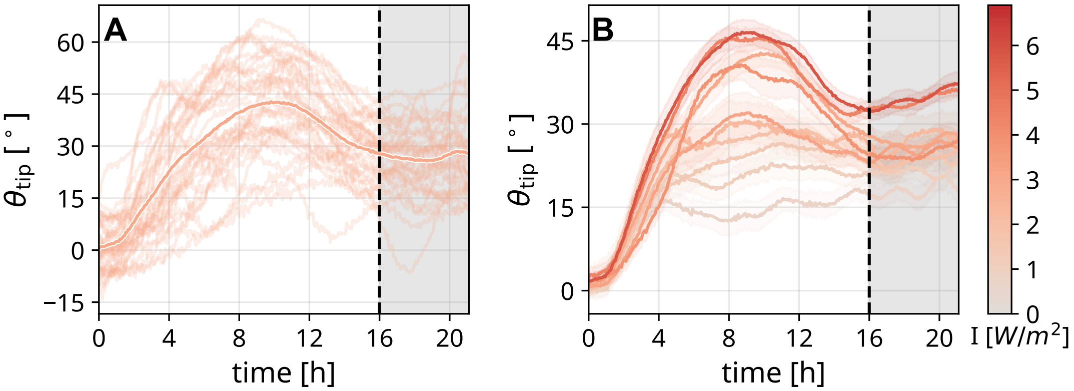


**Fig. S2. Full experimental results (A) example results for single light intensity** trajectories of different plants subjected to light intensity of 3.0 $\mathrm{W/m^2}$, N=33. Mean trajectory is highlighted. **(B) all unilateral mean trajectories used.** We note that light intensities above $7.0$ $\mathrm{W/m^2}$ lead to a saturated tropic response.

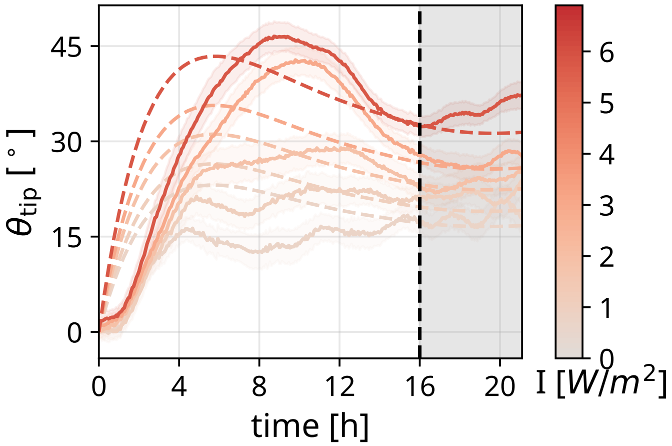


**Fig. S3. Comparison of simulated and experimental dynamics** mean tip angle trajectory over time for unilateral experiments plotted along with experimental results.

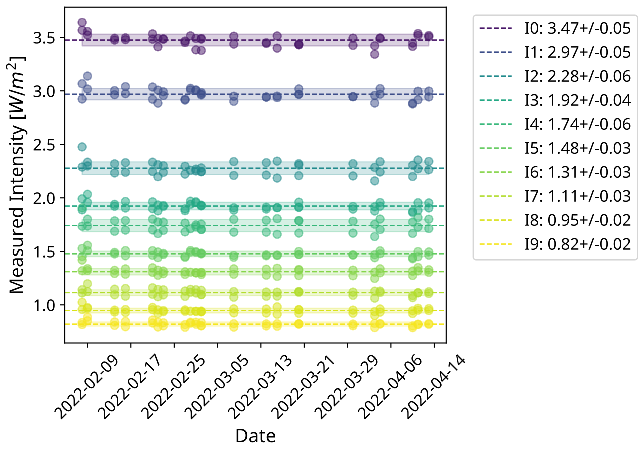


**Fig. S4. Estimation of error of light intensities** Light intensities were measured on multiple days to estimate the measurement uncertainty $dI$. Ten different intensities where sampled over 21 different days, and mean and standard deviation where calculated for each. We use the maximal STD value that was found, $dI = 0.06$ W/m$^2$.

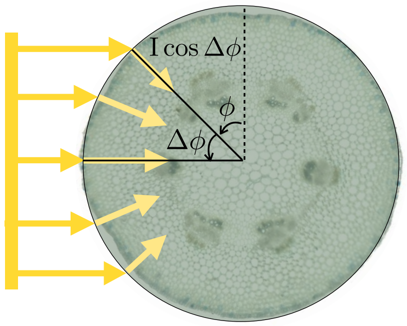


**Fig. S5. Angular attenuation across a cylindrical sensing surface.** Local light intensity depends on the cosine of the angle between the incident direction and the surface normal (Lambert's cosine law), leading to spatially nonuniform encoding around the circumference.

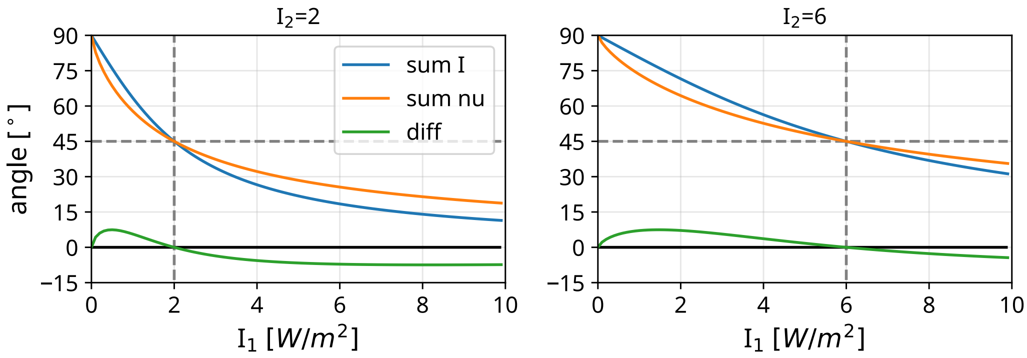


**Fig. S6. Specific examples of 2 perpendicular light sources** The difference between the direction of physical net incidence light, and the transduced signal direction for examples of light intensities set for one of the intensities and a sweep over the other one. We can see that when the light intensities are equal (denoted by the grey dashed line) there is no difference between the perceived and physical sum, and both predict a 45 degree angle.

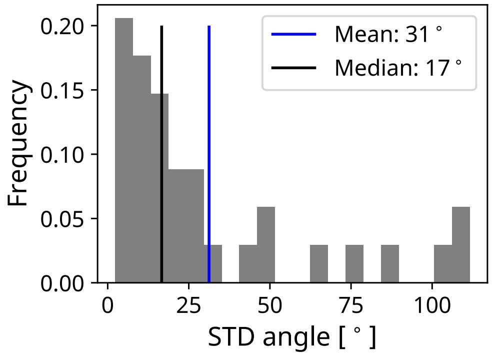


**Fig. S7. Inherent noise due to circumnutations.** For each plant in the out-of-plane illumination experiment (Fig. 4A, see sample trajectories in Fig. S8) we calculated the angle of the tip (top view) during the final 3 h. For each experiment we calculated the standard deviation of the angles, and plot here the distribution, including mean and median values.

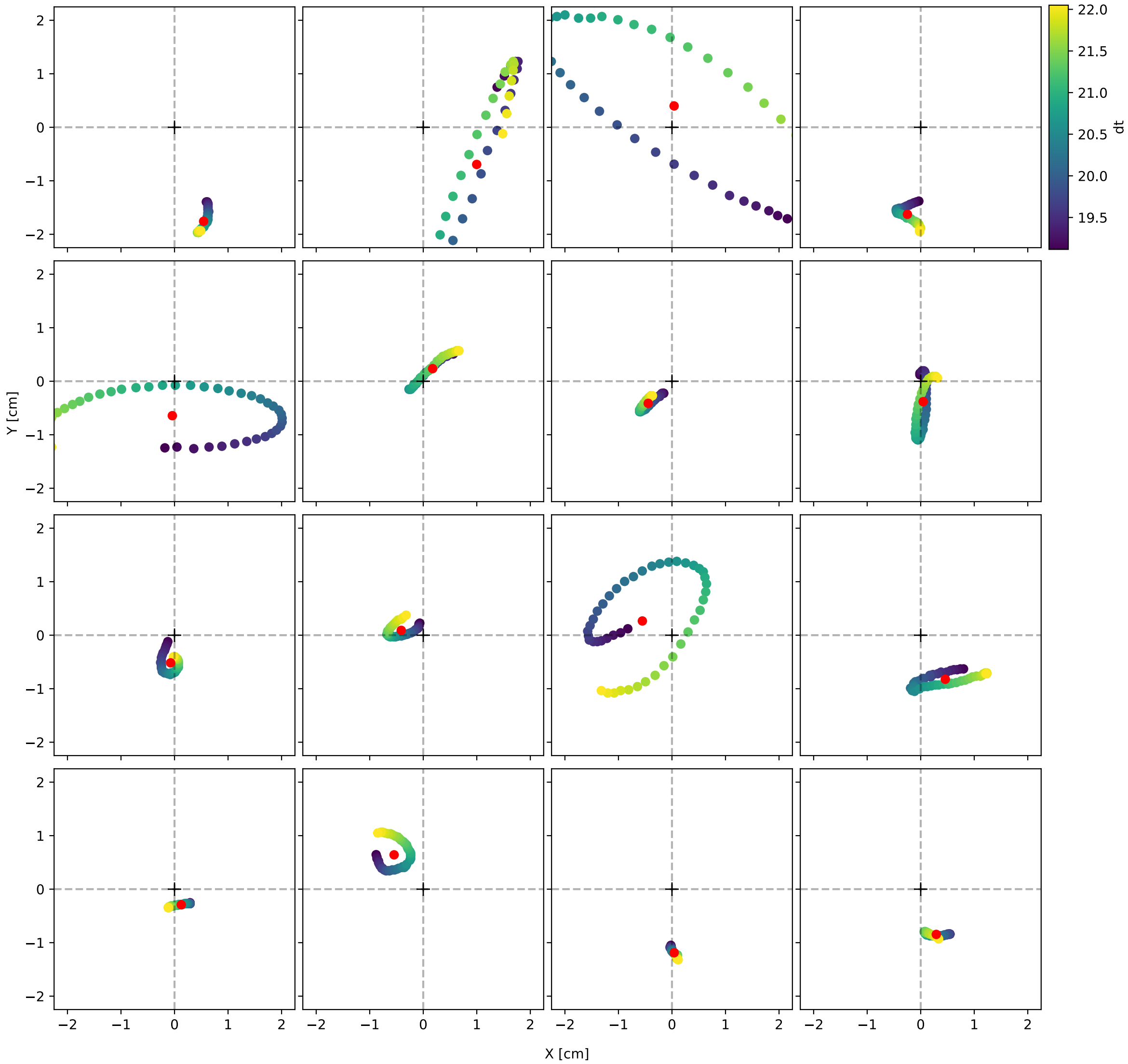


**Fig. S8. Sample top view trajectories** of plants grown with 3 light sources (Fig. 4A, exhibiting inherent movements termed circumnutations.

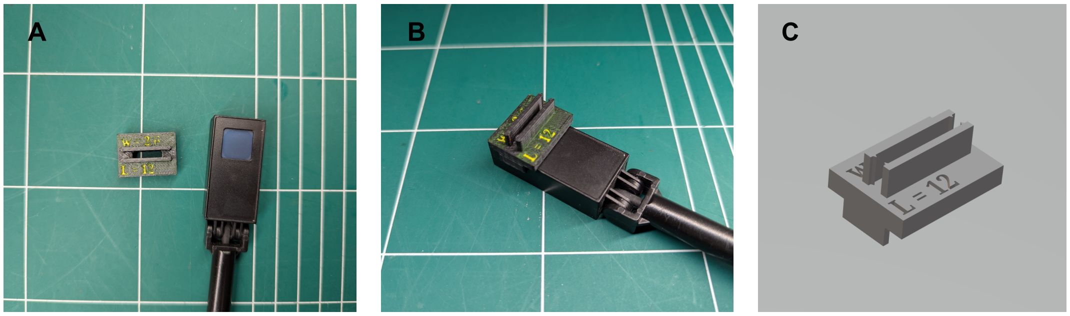


**Fig. S9. Measuring device for T.** A 3D printed sleeve was placed over the ophir sensor to measure the light intensity with and without a plant blocking the slit. **(A-B) Images of the device** The printed device (A) placed next to the sensor and (B) mounted onto the sensor. **(C) The .stl file for this device is can be found in the data repository.**

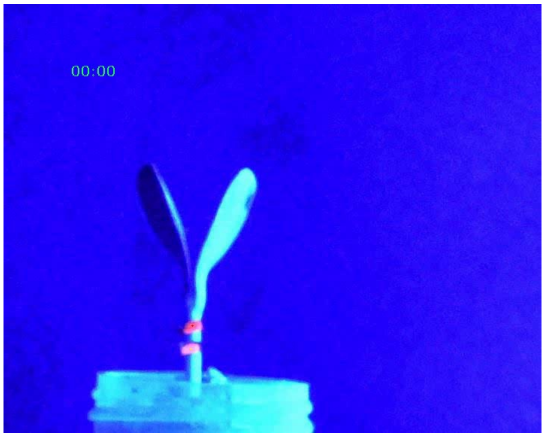


**Supplementary Video S1. Example of a tropic response.** Example of a single plant during the experiment with unilateral lighting. The fluorescent markers are used to for tracking purposes, and the trajectory of the tracking algorithm for both top and bottom points. Time (upper left corner) is shown in HH:MM format, counting from the beginning of the experiment (i.e. when the lights were turned on).
